# Alignment-free microbiome-based classification of fresh produce safety and quality

**DOI:** 10.1101/2022.08.25.505309

**Authors:** Chao Liao, Luxin Wang, Gerald Quon

**Author notes:** Co-corresponding authors: Luxin Wang, Associate Professor, University of California Davis, Davis, CA 95616,; Gerald Quon, Assistant Professor, University of California Davis, CA 95616,.

## Abstract

Small samples sizes and loss of up to 50-70% of sequencing data during the data denoising step of preprocessing can limit the statistical power of fresh produce microbiome analyses and prevent detection of important bacterial species associated with produce contamination or quality reduction. Here, we explored an alignment-free analysis strategy using k-mer hashes to identify DNA signatures predictive of produce safety and produce quality, and compared it against the amplicon sequence variant (ASV) strategy that uses a typical denoising step. Random forests (RF)-based classifiers for fresh produce safety and quality using 7-mer hash datasets had significantly higher classification accuracy than those using the ASV datasets. We also demonstrated that the proposed combination of integrating multiple datasets and leveraging an alignment-free 7-mer hash strategy leads to better classification performance for fresh produce safety and quality. Results generated from this study lay the foundation for future studies that wish and need to incorporate and/or compare different microbiome sequencing datasets for the application of machine learning in the area of microbial safety and quality of food.

---

Next generation sequencing approaches for the analysis of microbial communities in fresh produce include amplicon-based sequencing (e.g. 16S rRNA gene sequencing) and metagenomic sequencing (e.g. shotgun sequencing). These technologies help identify microbial populations within individual produce samples, traditionally by aligning sequenced reads to known genomes to quantify the presence of known species in a sample^1^. When multiple samples are sequenced, statistical and machine learning approaches can be leveraged to identify species of importance to produce safety (PS) and produce quality (PQ)^2–12^. Interactions between pathogenic and/or spoilage microorganisms and other native or background microbiota shine light on new competitive exclusion microorganisms to protect and improve the safety and quality of fresh produce^9,11–14^.

Of particular interest in this study is the use of machine learning to identify biomarkers: bacterial species whose presence is correlated with produce safety or quality. Biomarkers can be identified by constructing classification models that identify produce sample features that distinguish samples of different PQ or PS statuses. Both the accuracy of the classifiers, as well as the quality and reproducibility of the biomarkers identified from them, typically increase with dataset and sample sizes^15^. When reviewing published produce microbiome datasets, we found no overlap of PS-related biomarkers across three PS-related studies^11,12,16^, and only one biomarker consistently observed across three PQ-related studies^9,11,16^. This poor overlap of biomarkers suggests an opportunity to identify alternative biomarker identification strategies that identify more reproducible biomarkers across studies.

The poor overlap of biomarkers identified by different studies may be driven by two reasons. The first reason is due to the limited sample sizes in each individual study. On average, 110 samples are analyzed in previously published studies^1,4–12,17^, which is much smaller than that suggested given the large number of microbial species being identified in each sample^18^. Small sample sizes coupled with the profiling of many bacterial species can lead to more spurious (false) biomarkers, or correlations between bacterial species occurrence and pathogen contamination or quality decline, and ultimately yield poor reproducibility between studies^11–13^. Secondly, this poor overlap might be caused by low effective sequencing depth of each produce sample. Two key steps of microbiome sequencing data analysis are the alignment of reads to known genomes used in the metagenomic sequencing analysis, and the denoising step used in the amplicon sequencing analysis. During this process, up to 70% of reads for the metagenomic sequencing and 50% of reads for the 16S rRNA gene sequencing can be removed or “wasted” due to their poor matching to the database and nosing sequences^19,20^. This loss of over half of the sequencing reads reduces the power to identify low abundant species or sequence variations^21^.

Here we proposed a computational strategy to address both challenges in order to better identify bacterial biomarkers broadly correlated with both PS and PQ phenotypes. First, we used an alignment-free approach based on counting short k-mers in sequencing reads^22,23^ to characterize microbiota in individual fresh produce samples (**Fig. 1A**). We showed that k-mer hash-based models were significantly better at predicting both PS and PQ compared to the commonly used approach of counting amplicon sequence variants (ASV). Second, we integrated PS and PQ datasets before analysis to boost dataset size and power, and show by doing so we are able to identify taxa that are more broadly associated with PS and PQ.

**Fig. 1.**
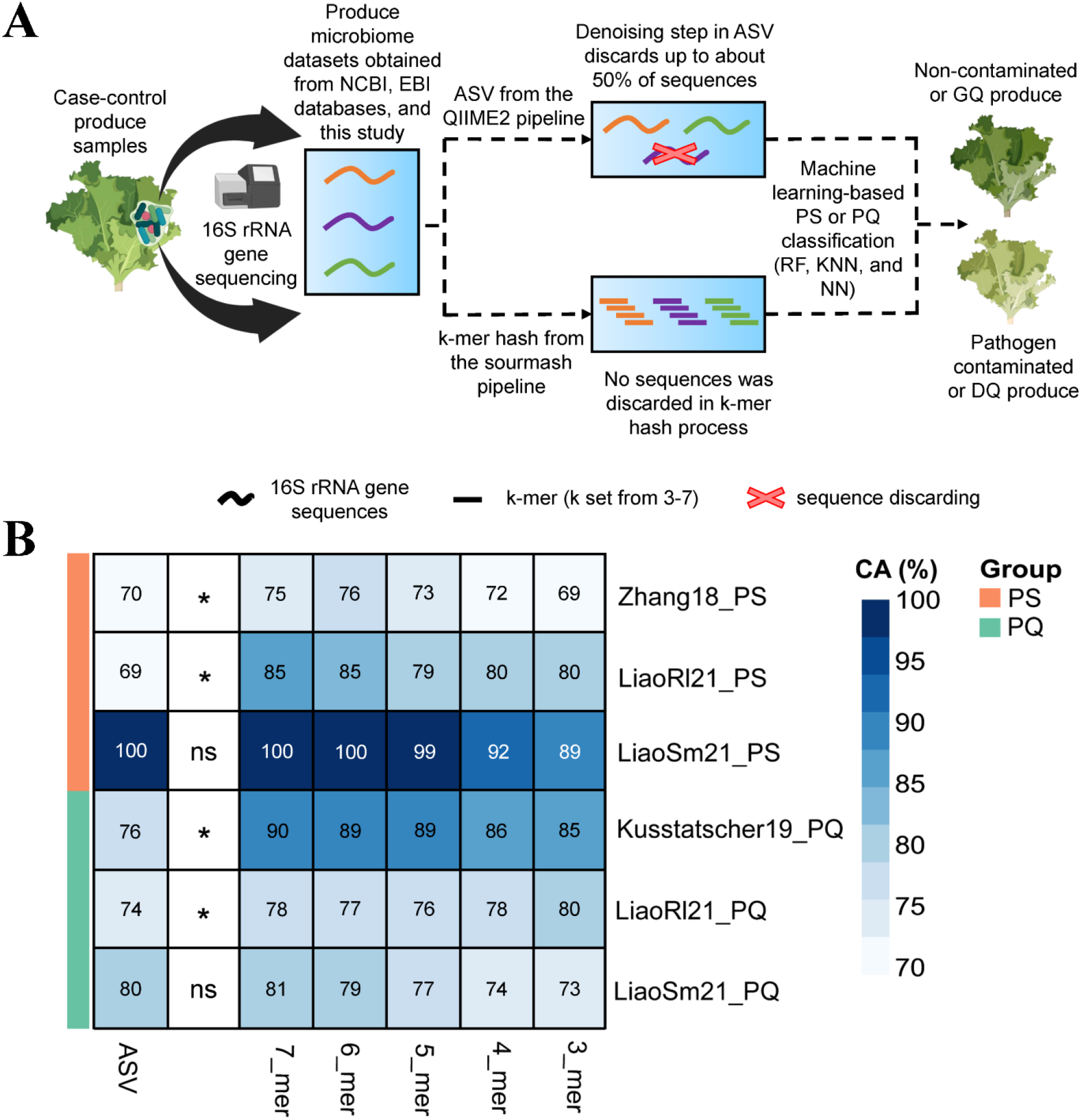
(**A**) Schematic comparing the preprocessing including denoising of the ASV approach against the alignment-free preprocessing of the k-mer hash approach for constructing produce safety (PS) and produce quality (PQ) classifiers. (**B**) Heatmap of the accuracies achieved by RF-based models using fresh produce microbiome datasets associated with PS and PQ in ASV format and k-mer hash format (k from 3 to 7). Accuracies are measured using 10-fold cross-validation. GQ and DQ represent good-quality produce and decreasing-quality produce, respectively. Partial icons in (A) were obtained from “BioRender.com”.

## Results

### Alignment-free analysis leads to more accurate classification of microbiome samples

We first tested the hypothesis that using a k-mer hash strategy to analyze 16S rRNA sequencing data that avoids read alignment and the corresponding data loss in the ASV strategy would improve PQ and PS classification accuracy and biomarker identification. **Fig. 1A** illustrates the conceptual differences between the two strategies. For each individual PS and PQ study, we constructed machine learning-based classifiers by selecting one of three methods (RF, *k*-NN, and NN), and training them using either the ASV or k-mer hash representations of the study data. Overall, the RF-based classifiers using 7-mer hash (**Fig. 1B**) representations showed higher classification accuracy than *k*-NN-based and NN-based for both PS (**Supplementary Fig. 1**) and PQ (**Supplementary Fig. 2**). RF-based classification using the 7-mer hash representation also generally outperformed classification using the ASV representation by 8% on average (*P* < 0.05, Wilcox rank sum test), except for the LiaoSm21_PS and LiaoSm21_PQ studies. In LiaoSm21_PS both the ASV and 7-mer hash versions of the data yielded an accuracy of 100%. We also found the k-mer hash datasets performed better with larger k values within the range of 3 to 7. This is unsurprising because larger k values lead to more features extracted from the sequencing data, which in turn provides more opportunities to distinguish DNA sequence signatures between cases and controls.

### Integrated microbiome data analysis leads to more generalizable classification

To increase the effective sample sizes of published microbiome studies and therefore identify taxa that are robustly associated with PS and PQ, a single, integrated produce safety (IPS) dataset and a single, integrated produce quality (IPQ) dataset were established by applying batch correction and merging studies within each category. We computed ASV and 7-mer hash representations of the IPQ and IPS datasets, and used them to construct RF-based models whose classification performance were compared against our previous models constructed on individual studies. Consistent with the analyses conducted on the individual studies, the integrated 7-mer hash representation achieves higher classification accuracy compared to ASV representation by 5% (*P* < 0.05) and 17% (*P* < 0.05) for the IPS and IPQ datasets, respectively (**Fig. 2A**), supporting the notion that the 7-mer hash dataset retains more actionable information in sequencing reads. **Fig. 2B** illustrates the performance differences separated by component study, and we see that the classification performance is systematically higher in samples from all studies, not just a select study. We also found that for the IPS dataset, when we replaced the binary labels with the pathogen-specific labels (*E. coli* O157, *L. monocytogenes*, and *Salmonella* Infantis), performance similarly was higher for 7-mer representations (85%) compared to ASV (78%) (**Supplementary Fig. 7**). The accuracy of the 7-mer hash IPS classifier was 82%, and when the IPS classifier was used to predict specific pathogens, it rose to 85% (**Supplementary Fig. 7**). Our IPS classifier therefore performs as well as the commonly used culture-dependent approach using selective agars that does not rely on sequencing^24^. Given the size and number of PS datasets will increase over time, we expect sequencing-base classifiers to outperform the culture-dependent approach in the future. For the IPQ classifiers that achieved 82% accuracy, there are no benchmarks to compare its performance against as this was the first time predictive models were used to evaluate the quality of fresh produce using microbiota.

**Fig. 2.**
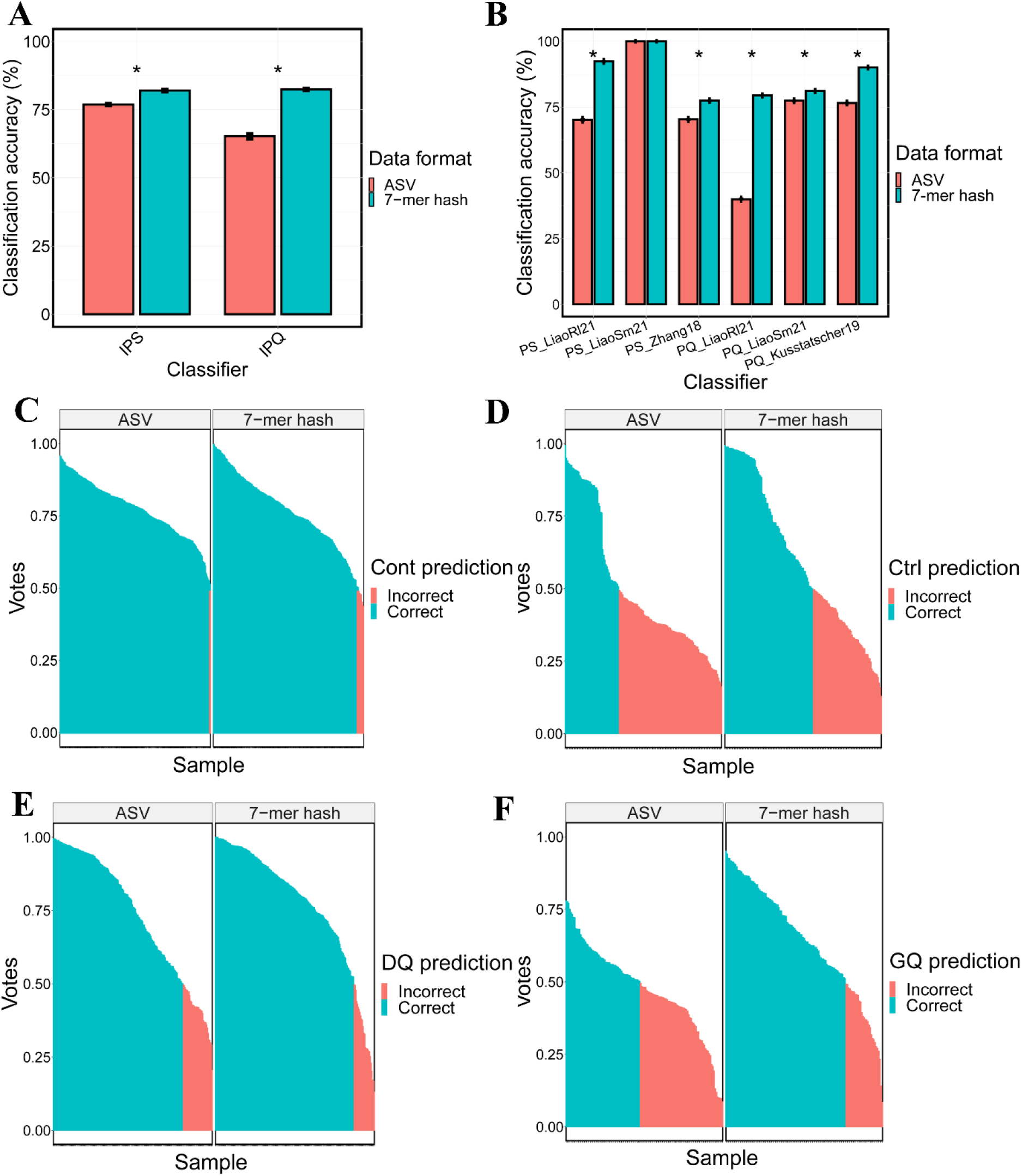
(**A**) Barplot of classification performance of RF-based models on the integrated PS (IPS) and integrated PQ (IPQ) datasets, represented using either ASV or 7-mer hashes. The Wilcoxon rank sum test was used for the pairwise comparison of the accuracies of the IPS and IPQ classifiers. (**B**) Barplot of classification performance of classifiers established by using individual datasets from IPS and IPQ datasets in ASV and 7-mer hash. (**C**) Barplot of the prediction of PS samples with the true label of contamination. The prediction was made by votes of 500 decision trees in RF-based classifiers established by using ASV and 7-mer hash. The cutoff voting rate (50% votes) indicates whether a labeled sample is predicted correctly or not. Ctrl and Cont represent non-contaminated samples and contaminated samples. The * stands for *P* < 0.05. (**D**) Same as (C), but barplot of the prediction of PS samples with the true label of control. (**E**) Same as (C), but barplot of the prediction of PQ samples with the true label of decreasing quality (DQ). (**F**) Same as (C), but barplot of the prediction of PQ samples with the true label of good quality (GQ).

To gain insight into which samples were better classified under the integrated 7-mer hash dataset, we then visualized the label predictions of individual samples generated by the RF-based classifiers. **Figs. 2C-2F** compares the voting rates based on 7-mer hash and ASV for predicted versus true labels of each sample in IPS and IPQ. For the IPS samples, while the number of correctly predicted contaminated samples is similar (254 and 264 for 7-mer hash and ASV, respectively) (**Fig. 2C**), there is a marked increase in accuracy for 7-mer hash predictions of the non-contaminated samples (85 and 52 for 7-mer hash and ASV, respectively) (**Fig. 2D**). Similarly, in the analysis of the IPQ samples, the 7-mer hash representation led to more accurate prediction of the decreasing-quality samples (174 and 163 for 7-mer hash and ASV, respectively) (**Fig. 2E**), while the number of correctly predicted “good-quality” samples was similar (100 versus 62 for 7-mer hash and ASV, respectively) (**Fig. 2F**). Our results support our hypothesis that the 7-mer hash representation leads to better classification performance of microbiome samples, consistent with our results on individual samples.

We next wondered to what extent integration of microbiome data from multiple studies explicitly led to the construction of more generalizable classifiers, compared to classifiers trained on individual studies. We therefore performed six experiments (three for each of IPS and IPQ), in which we repeatedly removed one study as a test (held-out) study, and compared the model performance of RF classifiers when trained on the remaining two studies separately versus combined (**Fig. 3**). In principle, classifiers trained on the combined datasets would be encouraged to learn taxa that are more broadly associated with either produce pathogen contamination or produce quality decline, because both the case and control samples in the combined dataset will be more heterogeneous compared to the samples in the individual studies. **Fig. 3** shows that across the six experiments, the integrated datasets performed significantly better than the individual datasets alone in three of them (**Figs. 3A**, **3B**, and **3D**), achieving on average 17% higher accuracy than individual datasets. In comparison, for only one experiment, combining data led to significantly worse performance (**Fig. 3C**). These results suggest that it can be sensible to combine data from multiple studies together, even if the phenotypes studied do not directly match up, and tend to lead to better generalizable performance of the classifiers for both PS and PQ.

**Fig. 3.**
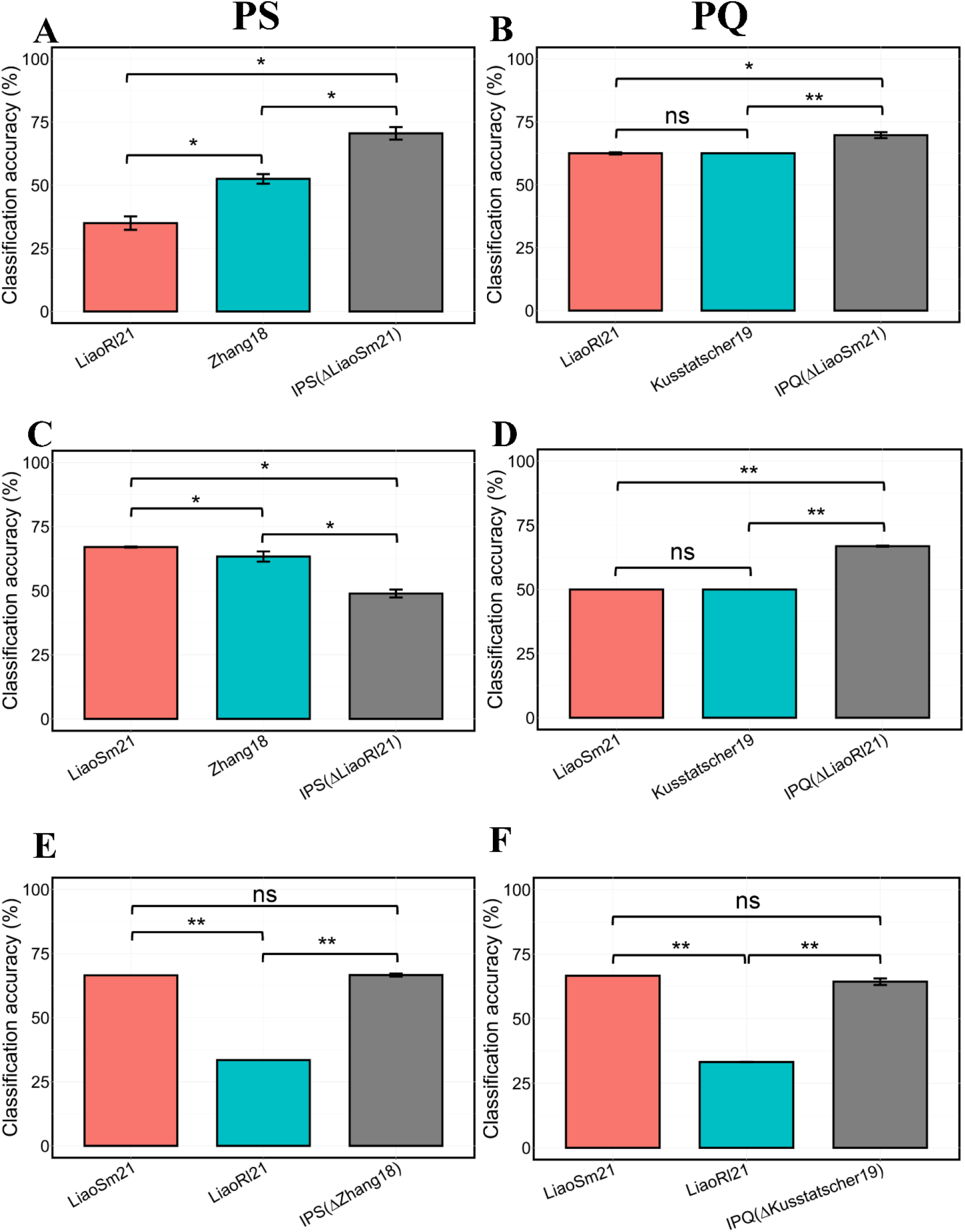
Barplots comparing the classification performance of classifiers trained on individual PS and PQ datasets, and IPS and IPQ datasets. (**A**) Classifiers were trained on individual LiaoRl21 or Zhang18 datasets and a classifier was trained on an integrated dataset excluding LiaoSm21, denoting IPS (ΔLiaoSm21). The LiaoSm21 was used as a testing set. (**B**) Classifiers were trained on individual LiaoRl21 and Kusstatscher datasets and an integrated dataset excluding LiaoSm21, denoting as IPQ (ΔLiaoSm21). The LiaoSm21 was used as a testing set. (**C**) Same as (B), but individual and integrated PS classifiers were tested on LiaoRl21. (**D**) Same as (B), but individual and integrated PQ classifiers were tested on LiaoRl21. (**E**) Same as (B), but individual and integrated PS classifiers were tested on Zhang18. (**F**) Same as (B), but individual and integrated PQ classifiers were tested on Kusstatscher19. Wilcoxon rank sum tests were applied for testing the significance of the difference in testing accuracy (%) of individual and integrated PS and PQ classifiers. PS and PQ mean produce safety and produce quality, respectively. The * represents *P* < 0.05; The ** stands for *P* < 0.01; The ns represents no significance.

### Integrated taxonomic analysis identifies generalizable taxa associated with pathogen contamination

The higher classification performance of the integrated datasets suggests that there are taxa that are broadly associated with PS and PQ identified by the classifiers. We therefore first inspected the 7-mer hash features identified as predictive by the classifier, to try to subsequently identify the underlying taxa contributing those predictive DNA signatures. Unfortunately, the current sourmash pipeline that was used to compute the k-mer hash signatures did not support mapping 7-mer hash signatures to the LCA database (see Methods). We therefore instead analyzed the classifiers trained on the ASV representation of the IPS and IPQ datasets for the following taxonomic analysis.

For the IPS taxonomic analysis, 1,131 genera were identified in total. The top 3 most dominant genera in the bacterial communities across fresh produce samples were *Pseudomonas* (with relative abundance ranging from 0.07% to 57.95%), unclassified genera (4.54% to 43.61%), and *Flavobacterium* (1.31% to 41.31%). *Pseudomonas* and *Flavobacterium* as dominant genera in bagged Spring mix salad and lettuce have been reported^11,25^. However, unclassified genera with high relative abundance were not often mentioned in fresh produce microbiota studies. The integration of datasets might bring out more unclassified genera than individual datasets, due to the limited number of taxonomic references available for taxonomic analysis.

The ANCOM-BC test was applied to identify bacteria at the genus level that have significantly different relative abundance between contamination groups (contaminated samples versus non-contaminated samples) or pathogen groups (*E. coli* O157, *L. monocytogenes*, and *Salmonella* Infantis). Five genera were identified as biomarkers for the contaminated group, including *Escherichia-Shigella, Listeria, Bacteroides, Peredibacter*, and *Faecalibacterium*, and two biomarkers (*Rheinheimera* and *Pseudomonas)* were identified for the non-contaminated group (**Fig. 4A**). Their significance values were listed in **Supplementary Table 3.** We noticed that the contaminating pathogens, *E. coli* O157 and *L. monocytogenes*, were accurately identified as *Escherichia-Shigella* and *Listeria*. Our previous study^11^ reported that no *Escherichia* could be identified in contaminated Spring mix samples when the *E. coli* O157:H7 inoculation level was at 5.5 Log CFU/ml by using 16S rRNA gene sequencing. The authors assumed that the concentration of *E. coli* O157 was below the limit of detection of 16S rRNA gene sequencing method. However, the present study showed the *Escherichia-Shigella* was identified for *E. coli* O157 with relative abundance from 0.019% to 0.18%. One explanation for the difference is that the present study conducted the taxonomic classification analysis based on the SILVA 16S rRNA sequence database, while the Liao and Wang (2021) applied Greengenes database, which contains less annotation taxonomic references than SILVA database^11^.

**Fig. 4.**
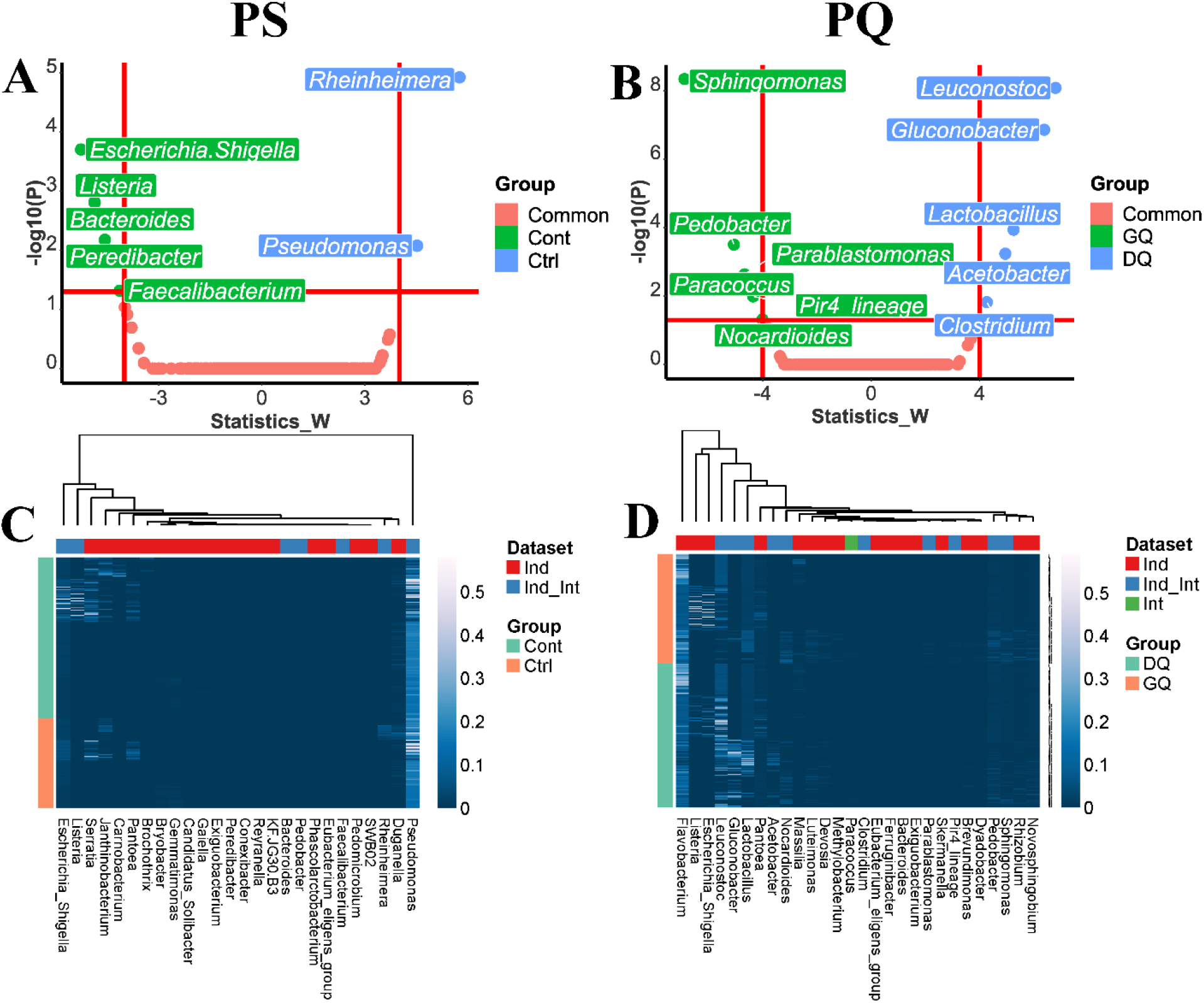
(**A**) Volcano plots based on the W statistic values and -log10(P) values obtained from the ANCOM-BC test presenting the differential abundances of genera among two contamination groups for PS. (B) Volcano plots based on the W statistic values and -log10(P) values obtained from the ANCOM-BC test presenting the differential abundances of genera among two quality groups for PQ. (**C**) Heatmap of the relative abundances of bacterial biomarkers identified from individual and integrated PS datasets. (**D**) Heatmap of the relative abundances of bacterial biomarkers identified from individual and integrated PQ datasets. Ind means individual datasets. Int represents the integrated dataset. Cont means contaminated samples. Ctrl represents non-contaminated samples. DQ represents decreasing-quality samples. GQ represents good-quality samples.

On the other hand, the *Salmonella* Infantis was not identified using the Zhang18 study (Zhang et al., 2018). One explanation is that 10^4^ copies/ml of *Salmonella* Infantis located in the root of lettuce was below the limit of detection of 16S rRNA gene sequencing. In this study, *Peredibacter* was identified as a biomarker for *Salmonella* Infantis-contaminated samples although *Salmonella* Infantis could not be detected. *Bacteroides* is an obligate anaerobic bacterium making up a remarkable portion of fecal bacterial communities, which has been suggested used as fecal indicator organisms for water samples^26^. Our result indicate that Bacteroides could be also applied as pathogen contamination indicator for produce. Davi-dov and Jurkevitch (2004) reported that *Peredibacter* is a member of *Bdellovibrio*-and-like organisms (BALO) that are highly motile microbes preying on other Gram-negative bacteria^27^. Lu and Cai (2010) reported that *Peredibacter* sp. strain BD2GS significantly retarded the growth of *Salmonella* Typhimurium within 3-12 hours through lysing prey cells^28^. Based on these, the elevated relative abundance of *Peredibacter* might be triggered by contamination of *Salmonella* Infantis. *Bacteroides* and *Faecalibacterium* have been reported as commensal bacteria of the human gastrointestinal microbiota^29,30^ and classified as fecal indicator bacteria^31^. Savichtcheva et al. (2007) reported that 16S rRNA gene marker of *Bacteroides* had a better prediction for the presence of bacterial enteric pathogens than total and fecal coliforms^32^.

### Integrated taxonomic analysis identifies generalizable taxa associated with produce quality

For the taxonomic analysis at genus level of integrated microbiome datasets related to PQ, 664 genera were identified. The top 3 genera based on the relative abundance were *Pseudomonas* (0.10% to 58.84%), *Flavobacterium* (0.73% to 38.53%), and Unclassified genera (5.49% to 55.52%). Through the ANCOM-BC test, five genera, including *Leuconostoc, Gluconobacter*, *Lactobacillus, Acetobacter*, and *Clostridium*, were identified as biomarkers in decreasing-quality samples (**Fig. 4B**). Their significance values were listed in **Supplementary Table 4.** Del Árbol et al. (2016) mentioned that *Gluconobacter* and *Acetobacter* are acetic acid bacteria, which can generate acetic acid to spoil fruits causing bacterial rot and browning^33^. *Leuconostoc* and *Lactobacillus* are commonly known as psychrotrophic spoilage lactic acid bacteria that spoil meat products and fresh fruits and vegetables during 4 °C storage^34,35^. *Clostridium* spp. have also been reported to be associated with meat and cheese spoilage^36,37^. In the Kusstatscher et al. (2019) study, except for *Clostridium*, the other four genera identified as biomarkers in decreasing-quality samples in the present study were recognized as core microbiota in decaying samples^9^. Six genera, including *Sphigomonas, Pedobacter, Parablastomonas, Paracoccus*, *Pir4_lineage*, and *Nocardioides*, were identified as biomarkers in good-quality samples. These six biomarkers for good-quality samples were reported to be related to agriculture rhizosphere soil^38–43^. Three genera, *Sphigomonas, Pedobacter*, and *Nocardioides*, were also identified as core microbiota in healthy samples in the Kusstatscher et al. (2019) study^9^.

Most of the biomarkers identified in the integrated datasets were covered by the biomarkers identified from individual datasets (**Supplementary Figs. 4** and **5**), but there were several new biomarkers identified solely in the integrated dataset. For the PS related datasets, seven biomarkers identified from the individual datasets (**Supplementary Fig. 4**) were also identified in the integrated dataset. However, seven biomarkers identified for the contaminated group in the individual datasets were not identified in the integrated dataset, including *Eubacterium*, *Janthinobacterium*, *Serratia*, *Carnobacterium*, *Phascolarctobacterium*, *Brochothrix*, and *Exiguobacterium*. Twelve biomarkers identified for the non-contaminated group in the individual datasets were not identified in the integrated dataset including *KF.JG30.B3, Gemmatimonas, Candidatus_Solibacter, SWB02, Conexibacter, Pedomicrobium, Bryobacter*, *Reyranella*, *Gaiella*, *Duganella*, *Pedobacter*, and *Pantoea*. (**Fig. 4C**).

For PQ datasets, 10 biomarkers identified in the integrated dataset were covered by the biomarkers identified in the individual datasets (**Supplementary Fig. 5**). Interestingly, *Paracoccus* was identified as a new biomarker for good-quality group only in the integrated dataset. To the best of our knowledge, *Paracoccus* has not been reported to be associated with produce quality. However, this genus contains a number of species that can produce Astaxanthin^44^, which has been reported to exhibit antagonism against food spoilage bacteria^45^. Fourteen bacteria identified as biomarkers for the good-quality group in the individual datasets were not identified in the integrated dataset, including *Flavobacterium*, *Ferruginibacter*, *Devosia*, *Skermanella*, *Luteimonas*, *Methylobacterium*, *Novosphingobium*, *Dyadobacter*, *Bacteroides*, *Exiguobacterium*, *Pantoea*, *Eubacterium*, *Escherichia*_*Shigella*, and *Listeria*. And four biomarkers recognized for decreasing-quality group in the individual datasets were not identified in the integrated dataset, including *Brevundimonas*, *Rhizobium*, *Pedobacter*, and *Dyadobacter* (**Fig. 4D**).

In addition to identifying critical features associated with PS and PQ using ANCOM-BC test, we ranked the features based on the MDA measures from RF-based PS and PQ classifiers established using individual and integrated ASV or 7-mer hash datasets and classified them into three groups, including negative, zero, and positive contributions to PS and PQ classification (**Fig. 5** and **Supplementary Fig. 3**). RF-based classifiers using the 7-mer hash representation utilized on average 62% of the 8192 hash features provided to RF for the integrated datasets, compared to an average of 9% of the ASV features provided to RF for the integrated datasets (**Supplementary Table 2**). Our interpretation is that the 7-mer hash representation may lead to better classification performance in part because so many features are leveraged in the classification; this may make classification more robust by making individual features weight less towards the label prediction.

**Fig. 5.**
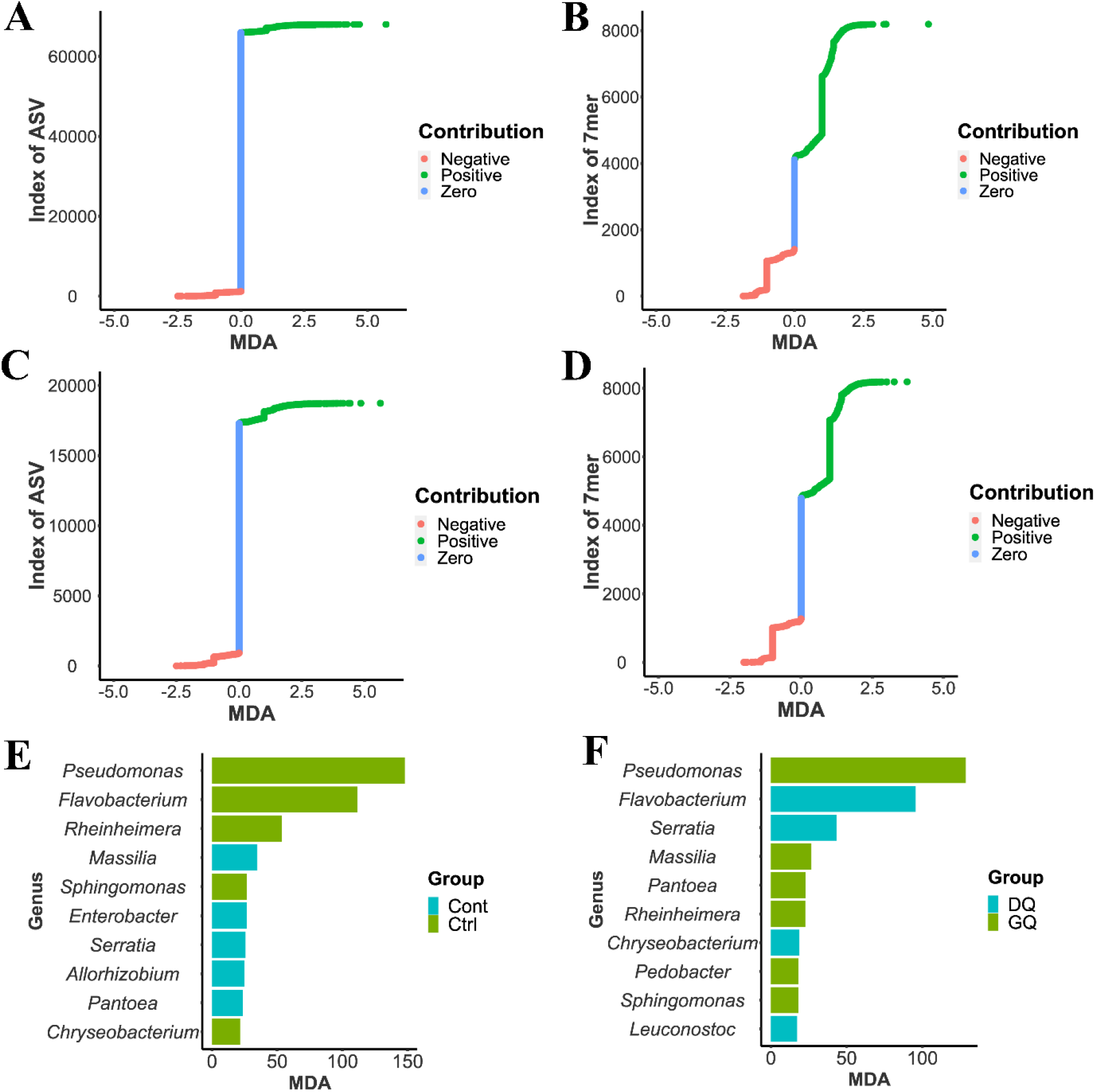
Importance of features evaluated by mean decrease accuracy (MDA) provided by random forests-based classifiers. (**A**) Contribution of ASV features to PS classification identified from RF-based IPS classifiers. (**B**) Contribution of 7-mer hash features to PS classification identified from RF-based IPS classifiers. (**C**) Same as (A), but for PQ. (**D**) Same as (B), but for PQ. (**E**) Identified top 10 genera with a positive contribution to IPS classification. (**F**) Identified top 10 genera with a positive contribution to IPQ classification. Ctrl represents the non-contaminated group and Cont represents the contaminated group. GQ represents good quality and DQ represents decreasing quality.

Overall, 332 genera were identified from the 2057 ASVs with a positive contribution to the IPS classification. The top 10 most important genera were *Pseudomonas*, *Flavobacterium*, *Rheinheimera*, *Massilia*, *Sphingomonas*, *Enterobacter*, *Serratia*, *Allorhizobium*, *Pantoea*, and *Chryseobacterium* (**Fig. 5E**). *Pseudomonas*, *Massilia*, *Enterobacter*, *Serratia*, *Allorhizobium*, and *Pantoea* were considered as the biomarkers for contaminated samples. Our previous study^11^ has reported that *Massilia* was identified as a biomarker for *E. coli* O157 contamination of Spring mix salad. Oliveira et al. (2015) have reported that *Enterobacter, Serratia*, and *Pantoea* isolated from lettuce presented an inhibitory effect against *Salmonella* and *L. monocytogenes*^46^. *Allorhizobium* was not reported previously to be associated with foodborne pathogen contamination of fresh produce, but some species are plant pathogens, e.g. *A. vitis* can infect grapevines^47^. Similarly, 138 genera were identified from the 1451 ASVs with positive contributions to the IPQ classification. Among them, the top 10 genera were *Pseudomonas, Flavobacterium*, *Serratia*, *Massilia*, *Pantoea*, *Rheinheimera*, *Chryseobacterium*, *Pedobacter*, *Sphingomonas*, and *Leuconostoc* (**Fig. 5F**). *Flavobacterium, Serratia, Chryseobacterium*, and *Leuconostoc* were regarded as biomarkers for the decreasing-quality samples. *Flavobacterium* has been reported to be associated with the spoilage of meat, milk, and seafood^48^. Hernández et al. (2020) reported *Flavobacterium* and *Chryseobacterium* belong to the class Flavobacteriales were dominant during the ready-to-eat lettuce spoilage but their importance during vegetable spoilage was still unclear^25^. Our results filled the knowledge gap to present the importance of *Flavobacterium* and *Chryseobacterium* during the quality reduction of fresh produce. In addition, it has been widely known that *Serratia* and *Leuconostoc* are microorganism that can cause food spoilage^48^. In addition to these four biomarkers, the RF-selected features contained another 41 biomarkers for decreasing-quality produce.

## Discussion

The integrated PS and PQ classifiers using 7-mer hash datasets had significantly higher accuracy than the models using ASV datasets. Two reasons are proposed: first, to obtain the ASV representations, the DADA2 plugin in QIIME 2 was used to process the primer-removed sequences, including quality filtering, denoising, chimera removing, dereplicating, and pair-end sequence joining. Although the parameter “--p-trunc-len” was set as 0 to trim no base due to the median quality score greater than 30 in this study, 1.66% to 40.94% of sequences from the integrated dataset related to PS were still discarded (**Supplementary Fig. 6A**) and 0.50% to 46.98% of overall sequences from integrated dataset related to PQ were discarded (**Supplementary Fig. 6B**). For the k-mer hash representations, the raw sequences were directly used to generate the k-mer hash datasets by using the sourmash pipeline as the all the median phred quality scores of nucleotides in reads were greater than 30^49^. The discarded reads may contain abundant sequence variation information, which contributed to the downstream sample classification. Werner et al. (2012) reported that 43% of total 16S rRNA gene sequences were discarded after the denoising process^21^. Second, the 7-mer hash computed from the original reads could improve the detection sensitivity of bases than ASV in a longer size (600 bp). The raw sequences were sub-sequenced into seven-base subsequences (7-mer), then transformed into hash codes, and finally randomly picked for 8,192 of 7-mer hash signatures for each sample^22^. The 7-mer hash datasets for establishing classification models can significantly shorten the computing time and improve the specificity and sensitivity of detection of different nucleotides among sequences compared with ASV.

The 3-mer to 7-mer hash datasets of individual studies related to PS and PQ were computed by using the sourmash pipeline, which were used subsequently to establish PS and PQ classifiers. Generally, the RF-based models using 6-mer and 7-mer hash datasets had similar accuracy but significantly higher accuracy than that of models using 3-mer to 5-mer hash datasets, except for PQ models using 3-mer of LiaoRl21. Sourmash pipeline generates k-mer minhash signatures of DNA sequences, which randomly samples k-mer content to produce small subsets as known as “sketches”^22^. The Jaccard similarity of two sketches of sequence data sets remains approximately equal to their true Jaccard similarity^50^. The factor “scaled” was set as 1 in ‘sourmash sketch dna’ command to generate the hash signature without down sampling the number of sketches. Using ‘sourmash sketch dna’ command with the k-mer size parameter set as 3 to 7 generated 32, 136, 512, 2080, and 8192 k-mer hashes of each DNA sample, respectively. Based on this, 7-mer hashes with a larger size than 3-mer to 6-mer contain more sequence variances, which contributes to the higher accuracy of models^51^. In addition, a larger k size of k-mer improves the specificity of different bases among sequences^52^. However, when k was set greater than 7, e.g. when k = 8, there were 32,896 of 8-mer hashes computed. The computing time and storage space remarkably increased while the accuracy of models had no obvious improvement (data not shown).

In the previous PS and PQ studies, a few bacterial biomarkers for produce contamination^11^ or decaying produce^9^ have been identified. However, these biomarkers were not consistent across the individual datasets due to the limited data size. In this study, we applied the data integration method to homogenize three PS datasets and three PQ datasets, respectively. The ANCOM-BC test was then employed to identify biomarkers for pathogen contamination and quality reduction of fresh produce based on individual datasets and integrated datasets. We identified 26 genus biomarkers and 28 genus biomarkers for PS and PQ produce, respectively, from individual datasets. Among them, seven biomarkers related to PS and 10 biomarkers related to PQ were validated by ANCOM-BC test using the integrated datasets. These validated biomarkers can provide a more generalizable and consistent indication or prediction of PS or PQ statuses^53^. Interestingly, we also identified a new biomarker, *Paracoccus*, for good-quality produce only from the integrated PQ dataset. The result indicates that more new biomarkers could be potentially identified with the size increase of integrated datasets. The non-validated biomarkers from individual datasets may be due to these studies having different types of fresh produce, distinct inoculated pathogens, and/or various storage conditions, which make the composition and diversity of bacterial communities largely different.

In addition to the ANCOM-BC test, the RF feature selection method was also used to identify biomarkers for fresh produce contamination and quality reduction. Compared to the biomarkers identified by the ANCOM-BC test, we found RF-based feature selection method can identify much more biomarkers for PS (332 genera) and PQ (138 genera) with a positive contribution to the PS and PQ classification. samples. These features covered all the biomarkers identified by ANCOM-BC test, indicating the RF-based models are more sensitive and powerful to catch the variation of features between classification groups. RF-based model and ANCOM-BC test can be used together to determine the reliable biomarkers for indicating PS and PQ. Sheh et al. (2022) reported that RF-based models were the most accurate models and correctly classified strictures indicators for chronic gastrointestinal diseases using 9 ASVs^54^. Based on the RF-based model and ANCOM results, *Clostridium perfringens* was identified as a potential causative agent associated with the development of strictures.

## Conclusion

In summary, we established and compared RF-based PS and PQ classifiers by using publicly available microbiome datasets in ASV and 7-mer hash representations for predicting the contamination conditions or quality statuses. This study illustrates that the alignment-free 7-mer hash-based approaches are useful for building more accurate classifiers than ASV, but not necessarily for the taxonomic analysis yet. Data integration of multiple datasets leads to greater classification performance of the integrated RF-based models than that using individual datasets, with significantly higher accuracy and more features with positive contribution to PS or PQ classification identified. In addition, we found more consistent and generalizable microbes were identified as biomarkers for safety and quality groups of fresh produce through integrated taxonomic analysis, illustrating the benefits of integrating datasets.

## Methods

### Fresh produce microbiome datasets

Four published fresh produce microbiome studies that generated 16S rRNA gene sequencing were selected, as they all have complete metadata information and are associated with PS and/or PQ. These four studies are named Zhang18^12^ (n = 236), Kusstatscher19^9^ (n = 240), LiaoSm21^11^ (n = 72), and LiaoR121 (n = 108) from the project LiaoRl21^16^ (this study) (**Table 1**). Project LiaoRl21 consists of combining published non-contaminated samples (n = 36)^16^, with unpublished data from pathogen-contaminated samples (n = 72) that were prepared and sequenced for this paper. The preparation of the contaminated samples, DNA extraction step, as well as library preparation and sequencing followed the protocol published by Liao and Wang (2021)^11^. The new data has also been uploaded in the public database. Based on these four studies, we created the PS dataset composed of case-control studies in which fresh produce was artificially inoculated with *Escherichia coli* O157:H7 (LiaoSm21 and LiaoRl21), *Listeria monocytogenes* (LiaoRl21), or *Salmonella* Infantis (Zhang18). Produce samples without contamination of pathogens were labeled as noncontaminated, while all other samples were given the uniform label contaminated. Also from these four studies, we created the PQ dataset in which samples were either labeled as good-quality (GQ, samples sequenced before their use-by dates or showing no decaying signs) or decreasing-quality (DQ, samples after their use-by date or decayed) (LiaoSm21, LiaoRl21, and Kusstatscher19).

**Table 1.**
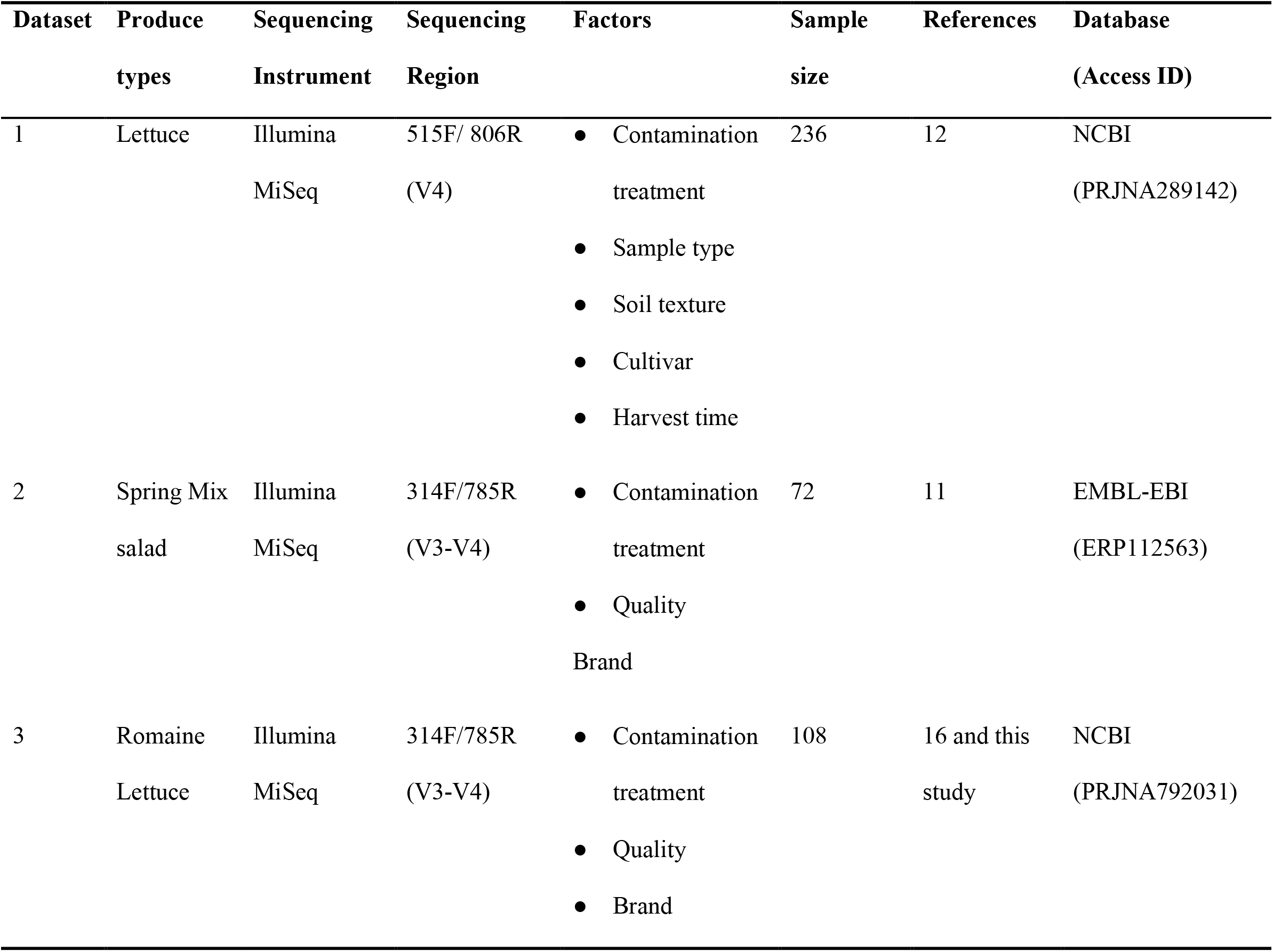
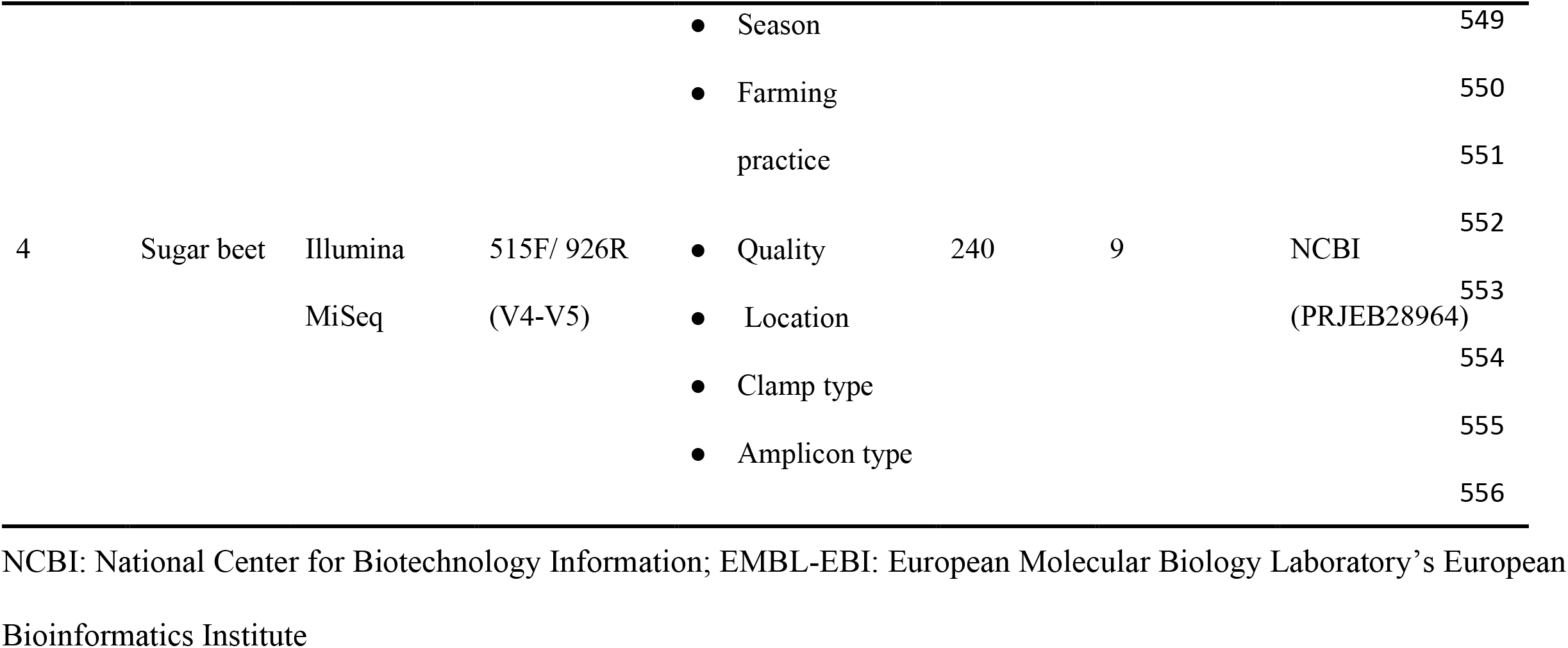
Characteristics of fresh produce microbiome datasets

### ASV dataset and k-mer hash dataset preparation

For each dataset, raw 16S rRNA gene sequences were imported into the QIIME 2 (Version 2021.8) pipeline^55^ by using the “qiime tools import” command. The barcodes and primers were removed by using “qiime cutadapt trim-paired/single” plugin of cutadapt^56^. Reads with median Phred quality scores of less than 30 were removed by truncating reads with a certain length using the “qiime dada2 denoise-paired/single” plugin of DADA2^49,57^. During this process, raw sequences were filtered, denoised, dereplicated, and clustered into ASV^58^. Sequences with barcode and primer sequences removed were then processed for quality control by truncating bases with the median Phred quality scores of less than 30 with the DADA2 package in R without ASV clustering^49,57^. After that, the processed sequences were used to compute k-mer hash signatures (k=3,…,7) for each sample by using the “sourmash sketch dna” command in the Sourmash pipeline^22^ (version 4.2.3)^22^.

### Common sum scaling

Since the sequencing depth varies across samples, datasets, and studies, the common sum scaling (COM) method was applied to normalize the sequencing depth among them for both ASV and k-mer hash methods. The ASV and k-mer hash counts were scaled to the minimum depth of each sample with the following equation^59^:

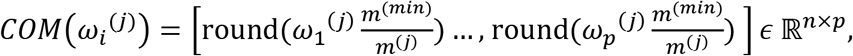

where *i* = 1,…,*p* is the ASV or k-mer index; *j* = 1,…, *n* is the sample index; 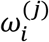 is the read count of ASV or k-mer *i* in sample *j*; *m*^(*j*)^ is the total ASV or *k*-mer hash count number for sample *j*, where 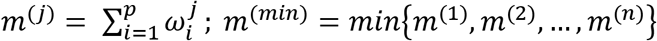 and round() is an operator rounding the fraction to be the nearest integer.

### Data integration and the removal of confounding factors

Data integration by using the Combat function in the ‘sva’ R package (version 3.42.0) was applied to remove batch effects indicated in the study’s metadata, such as sample type, location, and study-of-origin^60^. The parameter “par.prior” was set as “false” to use the nonparametric adjustments, and the parameter “mean.only” was set to “true” to adjust the mean of the batch effect across batches^60^.

### Classification of fresh produce safety and quality samples

Classifiers for PS and PQ using either the processed ASV or k-mer hash datasets were constructed. Three different classification methods, including random forests (RF)^61^, *k*-nearest neighbors (*k*-NN)^62^, and fully connected, feed-forward neural networks (NN)^63^ were evaluated (**Supplementary Figs. 1 and 2**). For RF, the number of decision tree (n_tree_) was set at 500, and the number of features randomly sampled as candidates at each split (m_try_) was set to the square root of the total number of input features. RF-based models were trained by using the “randomForest” R package^64^ (version 4.6-14). Predicted class labels were decided based on the majority vote (>50%) by 500 decision trees. The *k*-NN-based classifiers were trained by using the ‘caret’ R package (version 6.0-90). *k* values ranging from 1 to 10 were tested in order to identify *k* values with the best classification performance (**Supplementary Figs. 1 and 2**). The feed-forward NN were trained by using the ‘nnet’ R package (version 7.3-17), with hyperparameters set as the same as Arbajian et al. (2019)^65^ and Džal et al. (2021)^66^. The ‘nnet’ fits a feed-forward NN with a single hidden layer. The number of nodes in the hidden layer was set to 5, the decay parameter was set to 0.1, and the activation function was set to the logistic activation function.

### Model validation

For the individual dataset classification experiments, we used 10-fold cross validation to measure the accuracy of RF-, *k*-NN- and NN-based classifiers. Error was measured as the unweighted total classification accuracy across both the case and control phenotypes, as the classification experiments were generally well balanced (**Supplementary Table 1**). For RF-based models, the out-of-bag (OOB) estimate was also calculated in order to validate the models trained on PS or PQ datasets^61^.

A number of cross-study classification experiments were also conducted in order to test the generalization performance of PS and PQ classifiers. In the cross-study classification experiments for PS, we constructed pairs of training and test datasets in which one study formed the test dataset and the remaining studies formed the training dataset. For example, for the PS experiments, we constructed a training dataset consisting of the LiaoRl21 and Zhang18 studies, while using the LiaoSm21 study as a test dataset.

### Evaluation of feature importance

To evaluate and quantify the importance of individual features identified based on the ASV or the k-mer hash method, features were ranked by their mean decrease in accuracy (MDA) calculated by the random forests classification model. The MDA quantifies the importance of a variable by measuring the decrease in prediction accuracy when the variable is randomly permuted in comparison with the original observations^67^.

### Taxonomic analyses

To profile the bacterial communities within individual produce samples and carry out the differential abundance analysis between phenotypes, taxonomic analyses at the genus level were conducted by using the QIIME 2 (Version 2021.8) pipeline for the ASV dataset. The QIIME 2 plugin “q2-feature-classifier”^68^ was used to assign ASV to the SILVA 138 small subunit rRNA database as the taxonomy reference^69^. Chloroplast, mitochondria, and unassigned taxa were filtered by using “qiime taxa filter-seqs” command in QIIME 2^70^. For integrated taxonomic datasets, the COM method was first employed for data normalization. Data integration was carried out by using Combat to regress out study-of-origin, sample types, and storage conditions for PS and PQ datasets. The processed taxonomic datasets were employed for the differential abundance analysis between “contaminated” and “non-contaminated” samples, as well as between GQ and DQ samples.

For taxonomic analysis of 16S rRNA gene sequences based on the k-mer hash datasets, the SILVA 138 database was first downloaded and the taxonomic sequences were computed for 7-mer hash using the command “sourmash sketch dna” in the sourmash pipeline. The sourmash lowest common ancestor (LCA) taxonomy database was established by using a “sourmash lca index” command on the processed 7-mer hash of taxonomic sequences from the SILVA 138 database, followed by taxonomic classification of query 7-mer hash datasets against the established sourmash LCA taxonomy database carried out by using a command “sourmash lca classify”^22^. However, the query k-mer hashes could not be accurately assigned to the sourmash LCA taxonomy database based on SILVA database. This issue may be due to the inappropriateness of the current algorithm of LCA taxonomic classification in the sourmash pipeline for 16S rRNA gene sequences. This is an established issue (https://github.com/sourmash-bio/sourmash/issues/1421). Therefore, the taxonomic analysis here was performed only on the ASV representation of the data.

### Data visualization and statistical analysis

The non-parametric Wilcoxon rank sum test was applied to test for significant differences in classification accuracy between pairs of RF-based, *k*-NN-based, and NN-based classifiers. The analysis of composition of microbiome with bias correction (ANCOM-BC) test and the RF-based feature selection method mentioned above were employed to identify the bacterial biomarkers that had significantly abundance changes in one group over another^71^. All the above statistical analyses were conducted in R (version 4.1.0).

## Data availability

Raw reads from this study have been deposited to the National Center for Biotechnology Information under the project accession number PRJNA792031. The sample dataset is available in the GitHub repository (https://github.com/LZC0034/Nature-Food). Additional datasets that support the findings of this study are available from the corresponding authors upon request.

## Code availability

The codes for data processing are available in the GitHub repository (https://github.com/LZC0034/Nature-Food).

## Acknowledgements

This work was partially funded through a UC Davis CeDAR Innovative Data Science Seed

Funding Program Grant. G.Q. was supported by NSF CAREER award 1846559.

## Author contributions

C.L., L.W. and G.Q. conceptualized the study; C.L. performed the experiments, analyzed the data, and wrote the manuscript; L.W. and G.Q. supervised the study, reviewed and edited the manuscript.

## Additional information

Supplementary information. The online version contains supplementary material available at

## References

1. Jackson, C., Stone, B. & Tyler, H. Emerging perspectives on the natural microbiome of fresh produce vegetables. Agriculture 5, 170–187 (2015).

2. Bergholz, T. M., Moreno Switt, A. I. & Wiedmann, M. Omics approaches in food safety: fulfilling the promise? Trends Microbiol. 22, 275–281 (2014).

3. Ceuppens, S. et al. Characterization of the Bacterial Community Naturally Present on Commercially Grown Basil Leaves: Evaluation of Sample Preparation Prior to Culture-Independent Techniques. Int. J. Environ. Res. Public Health 12, 10171–10197 (2015).

4. Gu, G. et al. Shifts in spinach microbial communities after chlorine washing and storage at compliant and abusive temperatures. Food Microbiol 73, 73–84 (2018).

5. Jackson, C. R., Randolph, K. C., Osborn, S. L. & Tyler, H. L. Culture dependent and independent analysis of bacterial communities associated with commercial salad leaf vegetables. BMC Microbiol. 13, 274 (2013).

6. Jarvis, K. G. et al. Cilantro microbiome before and after nonselective pre-enrichment for Salmonella using 16S rRNA and metagenomic sequencing. BMC Microbiol. 15, 160 (2015).

7. Jarvis, K. G. et al. Microbiomes associated with foods from plant and animal sources. Front. Microbiol. 9, 2540 (2018).

8. Keshri, J. et al. Dynamics of bacterial communities in alfalfa and mung bean sprouts during refrigerated conditions. Food Microbiol 84, 103261 (2019).

9. Kusstatscher, P. et al. Microbiome-driven identification of microbial indicators for postharvest diseases of sugar beets. Microbiome 7, 112 (2019).

10. Leff, J. W. & Fierer, N. Bacterial communities associated with the surfaces of fresh fruits and vegetables. PLoS One 8, e59310 (2013).

11. Liao, C. & Wang, L. Evaluation of the bacterial populations present in Spring Mix salad and their impact on the behavior of Escherichia coli O157:H7. Food Control 107865 (2021). doi:10.1016/j.foodcont.2021.107865

12. Zhang, Y. et al. Microbial communities in the rhizosphere and the root of lettuce as affected by Salmonella-contaminated irrigation water. FEMS Microbiol. Ecol. 94, (2018).

13. Söderqvist, K. et al. Emerging microbiota during cold storage and temperature abuse of ready-to-eat salad. Infect Ecol Epidemiol 7, 1328963 (2017).

14. Yurgel, S. N., Abbey, Lord, Loomer, N., Gillis-Madden, R. & Mammoliti, M. Microbial Communities Associated with Storage Onion. Phytobiomes Journal 2, 35–41 (2018).

15. Costello, Z. & Martin, H. G. A machine learning approach to predict metabolic pathway dynamics from time-series multiomics data. NPJ Syst. Biol. Appl. 4, 19 (2018).

16. Liao, C. & Wang, L. The microbial quality of commercial chopped romaine lettuce before and after the “use by” date. Front. Microbiol. 13, 850720 (2022).

17. Dees, M. W., Lysøe, E., Nordskog, B. & Brurberg, M. B. Bacterial communities associated with surfaces of leafy greens: shift in composition and decrease in richness over time. Appl. Environ. Microbiol. 81, 1530–1539 (2015).

18. van der Ploeg, T., Austin, P. C. & Steyerberg, E. W. Modern modelling techniques are data hungry: a simulation study for predicting dichotomous endpoints. BMC Med. Res. Methodol. 14, 137 (2014).

19. Maran, M. I. J. & Davis G, D. J. Benefits of merging paired-end reads before pre-processing environmental metagenomics data. Mar. Genomics 61, 100914 (2022).

20. Prodan, A. et al. Comparing bioinformatic pipelines for microbial 16S rRNA amplicon sequencing. PLoS One 15, e0227434 (2020).

21. Werner, J. J., Zhou, D., Caporaso, J. G., Knight, R. & Angenent, L. T. Comparison of Illumina paired-end and single-direction sequencing for microbial 16S rRNA gene amplicon surveys. ISME J. 6, 1273–1276 (2012).

22. Pierce, N. T., Irber, L., Reiter, T., Brooks, P. & Brown, C. T. Large-scale sequence comparisons with sourmash. [version 1; peer review: 2 approved]. F1000Res. 8, 1006 (2019).

23. Vinje, H., Liland, K. H., Almøy, T. & Snipen, L. Comparing K-mer based methods for improved classification of 16S sequences. BMC Bioinformatics 16, 205 (2015).

24. Park, S.-H., Chang, P.-S., Ryu, S. & Kang, D.-H. Development of a novel selective and differential medium for the isolation of Listeria monocytogenes. Appl. Environ. Microbiol. 80, 1020–1025 (2014).

25. Hernandez, I. & Alfaro, B. Enhancing high throughput sequencing unveils changes in bacterial communities during ready-to-eat lettuce spoilage. Journal of Horticulture… (2020).

26. Tsai, K. et al. Bacteroides Microbial Source Tracking Markers Perform Poorly in Predicting Enterobacteriaceae and Enteric Pathogen Contamination of Cow Milk Products and Milk-Containing Infant Food. Front. Microbiol. 12, 778921 (2021).

27. Davidov, Y. & Jurkevitch, E. Diversity and evolution of *Bdellovibrio*-and-like organisms (BALOs), reclassification of *Bacteriovorax starrii* as *Peredibacter starrii* gen. nov., comb. nov., and description of the *Bacteriovorax-Peredibacter* clade as Bacteriovoracaceae fam. nov. Int. J. Syst. Evol. Microbiol. 54, 1439–1452 (2004).

28. Lu, F. & Cai, J. The protective effect of *Bdellovibrio*-and-like organisms (BALO) on tilapia fish fillets against Salmonella enterica ssp. enterica serovar Typhimurium. Lett. Appl. Microbiol. 51, 625–631 (2010).

29. Khan, M. T. et al. The gut anaerobe *Faecalibacterium prausnitzii* uses an extracellular electron shuttle to grow at oxic-anoxic interphases. ISME J. 6, 1578–1585 (2012).

30. Wexler, H. M. Bacteroides: the good, the bad, and the nitty-gritty. Clin. Microbiol. Rev. 20, 593–621 (2007).

31. Zhao, X.-L. et al. Coexistence of antibiotic resistance genes, fecal bacteria, and potential pathogens in anthropogenically impacted water. Environ. Sci. Pollut. Res. Int. 29, 46977–46990 (2022).

32. Savichtcheva, O., Okayama, N. & Okabe, S. Relationships between Bacteroides 16S rRNA genetic markers and presence of bacterial enteric pathogens and conventional fecal indicators. Water Res. 41, 3615–3628 (2007).

33. Toledo Del Árbol, J. et al. Microbial diversity in pitted sweet cherries (*Prunus avium* L.) as affected by High-Hydrostatic Pressure treatment. Food Res. Int. 89, 790–796 (2016).

34. Andreevskaya, M. et al. Food Spoilage-Associated *Leuconostoc, Lactococcus*, and Lactobacillus Species Display Different Survival Strategies in Response to Competition. Appl. Environ. Microbiol. 84, (2018).

35. Barth, M., Hankinson, T. R., Zhuang, H. & Breidt, F. in Compendium of the microbiological spoilage of foods and beverages (eds. Sperber, W. H. & Doyle, M. P.) 135–183 (Springer New York, 2009). doi:10.1007/978-1-4419-0826-1_6

36. Gómez-Torres, N. et al. Development of a specific fluorescent phage endolysin for in situ detection of Clostridium species associated with cheese spoilage. Microb Biotechnol 11, 332–345 (2018).

37. Palevich, N. et al. Comparative genomics of *Clostridium* species associated with vacuum-packed meat spoilage. Food Microbiol 95, 103687 (2021).

38. Asaf, S., Numan, M., Khan, A. L. & Al-Harrasi, A. Sphingomonas: from diversity and genomics to functional role in environmental remediation and plant growth. Crit Rev Biotechnol 40, 138–152 (2020).

39. Fagervold, S. K. et al. Microbial communities associated with the degradation of oak wood in the Blanes submarine canyon and its adjacent open slope (NW Mediterranean). Prog. Oceanogr. 118, 137–143 (2013).

40. Kwon, S.-W. et al. *Pedobacter rhizosphaerae* sp. nov. and *Pedobacter soli* sp. nov., isolated from rhizosphere soil of Chinese cabbage (*Brassica campestris*). Int. J. Syst. Evol. Microbiol. 61, 2874–2879 (2011).

41. Liu, X. et al. Biodegradation of aged polycyclic aromatic hydrocarbons in agricultural soil by Paracoccus sp. LXC combined with humic acid and spent mushroom substrate. J. Hazard. Mater. 379, 120820 (2019).

42. Shi, J. et al. Effects of wheat root exudates on bacterial communities in the rhizosphere of watermelon. Plant Soil Environ. 67, 721–728 (2021).

43. Takagi, K., Fujii, K., Yamazaki, K., Harada, N. & Iwasaki, A. Biodegradation of melamine and its hydroxy derivatives by a bacterial consortium containing a novel *Nocardioides* species. Appl. Microbiol. Biotechnol. 94, 1647–1656 (2012).

44. Fang, N. et al. De novo synthesis of astaxanthin: From organisms to genes. Trends Food Sci. Technol. 92, 162–171 (2019).

45. Mageswari, A., Subramanian, P., Srinivasan, R., Karthikeyan, S. & Gothandam, K. M. Astaxanthin from psychrotrophic *Sphingomonas faeni* exhibits antagonism against food-spoilage bacteria at low temperatures. Microbiol Res 179, 38–44 (2015).

46. Oliveira, M., Abadias, M., Colás-Medà, P., Usall, J. & Viñas, I. Biopreservative methods to control the growth of foodborne pathogens on fresh-cut lettuce. Int. J. Food Microbiol. 214, 4–11 (2015).

47. Habbadi, K. et al. Essential oils of Origanum compactum and Thymus vulgaris exert a protective effect against the phytopathogen Allorhizobium vitis. Environ. Sci. Pollut. Res. Int. 25, 29943–29952 (2018).

48. Odeyemi, O. A., Alegbeleye, O. O., Strateva, M. & Stratev, D. Understanding spoilage microbial community and spoilage mechanisms in foods of animal origin. Comp. Rev. Food Sci. Food Safety 19, 311–331 (2020).

49. Estaki, M. et al. QIIME 2 Enables Comprehensive End-to-End Analysis of Diverse Microbiome Data and Comparative Studies with Publicly Available Data. Curr Protoc Bioinformatics 70, e100 (2020).

50. Ondov, B. D. et al. Mash: fast genome and metagenome distance estimation using MinHash. Genome Biol. 17, 132 (2016).

51. Martínez-Porchas, M. & Vargas-Albores, F. Microbial metagenomics in aquaculture: a potential tool for a deeper insight into the activity. Rev. Aquacult. 9, 42–56 (2017).

52. Nasko, D. J., Koren, S., Phillippy, A. M. & Treangen, T. J. RefSeq database growth influences the accuracy of k-mer-based lowest common ancestor species identification. Genome Biol. 19, 165 (2018).

53. Kim, Y., Bismeijer, T., Zwart, W., Wessels, L. F. A. & Vis, D. J. Genomic data integration by WON-PARAFAC identifies interpretable factors for predicting drug-sensitivity in vivo. Nat. Commun. 10, 5034 (2019).

54. Sheh, A. et al. Alterations in common marmoset gut microbiome associated with duodenal strictures. Sci. Rep. 12, 5277 (2022).

55. Bolyen, E. et al. Reproducible, interactive, scalable and extensible microbiome data science using QIIME 2. Nat. Biotechnol. 37, 852–857 (2019).

56. Martin, M. Cutadapt removes adapter sequences from high-throughput sequencing reads. EMBnet j. 17, 10 (2011).

57. Callahan, B. J. et al. DADA2: High-resolution sample inference from Illumina amplicon data. Nat. Methods 13, 581–583 (2016).

58. Callahan, B. J., McMurdie, P. J. & Holmes, S. P. Exact sequence variants should replace operational taxonomic units in marker-gene data analysis. ISME J. 11, 2639–2643 (2017).

59. Badri, M., Kurtz, Z. D., Bonneau, R. & Müller, C. L. Shrinkage improves estimation of microbial associations under different normalization methods. NAR Genom. Bioinform. 2, lqaa100 (2020).

60. Leek, J. T., Johnson, W. E., Parker, H. S., Jaffe, A. E. & Storey, J. D. The sva package for removing batch effects and other unwanted variation in high-throughput experiments. Bioinformatics 28, 882–883 (2012).

61. Breiman, L. Random forests. Mach Learn (2001).

62. Altman, N. S. An Introduction to Kernel and Nearest-Neighbor Nonparametric Regression. Am. Stat. 46, 175–185 (1992).

63. Suykens, J. A. K. & Vandewalle, J. Least Squares Support Vector Machine Classifiers. Springer Science and Business Media LLC (1999). doi:10.1023/a:1018628609742

64. Liaw, A. & Wiener, M. Classification and regression by randomForest. R news (2002).

65. Arbajian, P., Hajja, A., Raś, Z. W. & Wieczorkowska, A. A. Effect of speech segment samples selection in stutter block detection and remediation. J. Intell. Inf. Syst. 1–24 (2019). doi:10.1007/s10844-019-00546-z

66. Džal, D. et al. Modelling bathing water quality using official monitoring data. Water (Basel) 13, 3005 (2021).

67. Calle, M. L. & Urrea, V. Letter to the editor: Stability of Random Forest importance measures. Brief. Bioinformatics 12, 86–89 (2011).

68. Bokulich, N. A. et al. Optimizing taxonomic classification of marker-gene amplicon sequences with QIIME 2’s q2-feature-classifier plugin. Microbiome 6, 90 (2018).

69. Quast, C. et al. The SILVA ribosomal RNA gene database project: improved data processing and web-based tools. Nucleic Acids Res. 41, D590–6 (2013).

70. McDonald, D. et al. An improved Greengenes taxonomy with explicit ranks for ecological and evolutionary analyses of bacteria and archaea. ISME J. 6, 610–618 (2012).

71. Lin, H. & Peddada, S. D. Analysis of compositions of microbiomes with bias correction. Nat. Commun. 11, 3514 (2020).

