## Supplementary Tables and Figures for "Alignment-free microbiome-based classification of fresh produce safety and quality"

**Supplementary Table 1.** The number of case and control samples in each fresh produce microbiome dataset

| Dataset | Class | Total sample number | Case number (Cont/DQ) | Control number (Ctrl/GQ) |
|---|---|---|---|---|
| Zhang18 | PS | 236 | 158 | 78 |
| LiaoSm21 | | 72 | 36 | 36 |
| LiaoRl21 | | 108 | 72 | 36 |
| Kusstatscher19 | PQ | 240 | 160 | 80 |
| LiaoSm21 | | 72 | 45 | 27 |
| LiaoRl21 | | 108 | 54 | 54 |

PS and PQ represent produce safety and produce quality; Cont and Ctrl stand for contaminated group and non-contaminated group; GQ and DQ mean good-quality group and decreasing-quality group.

**Supplementary Table 2.** The number of features with a positive or negative contribution to the PS and PQ classification based on mean decrease accuracy provided by RF-based models.

| Dataset | Class | Total number of ASV | Number of positive features (ASV) | Number of negative features (ASV) | Total number of 7-mer hash | Number of positive features (7-mer hash) | Number of negative features (7-mer hash) |
|---|---|---|---|---|---|---|---|
| Zhang18 | PS | 49404 | 2029 | 1253 | 8192 | 2665 | 1506 |
| LiaoSm21 | | 2510 | 401 | 46 | 8192 | 349 | 0 |
| LiaoRl21 | | 2365 | 228 | 276 | 8192 | 1114 | 543 |
| IPS | | 68007 | 2057 | 1179 | 8192 | 4084 | 1408 |
| Kusstatscher19 | PQ | 1164 | 210 | 66 | 8192 | 2070 | 603 |
| LiaoSm21 | | 2510 | 354 | 222 | 8192 | 908 | 448 |
| LiaoRl21 | | 2365 | 278 | 238 | 8192 | 1469 | 654 |
| IPQ | | 18732 | 1451 | 918 | 8192 | 3403 | 1271 |

PS and PQ represent produce safety and produce quality; IPS and IPQ stand for integrated produce safety and integrated produce quality. ASV means amplicon sequence variant.

**Supplementary Table 3.** *P* values of the biomarkers identified by the ANCOM-BC analysis for fresh produce in different contamination and pathogen groups.

| Biomarker | Group | *P* value |
|---|---|---|
| *Rheinheimera* | Ctrl (C) | $1.19 \times 10^{-5}$ |
| *Pseudomonas* | Ctrl (C) | $8.41 \times 10^{-3}$ |
| *Escherichia-Shigella* | Cont | $1.98 \times 10^{-4}$ |
| *Listeria* | Cont | $1.53 \times 10^{-3}$ |
| *Bacteroides* | Cont | $1.57 \times 10^{-3}$ |
| *Peredibacter* | Cont | $6.69 \times 10^{-3}$ |
| *Faecalibacterium* | Cont | 0.048 |
| *Rheinheimera* | Ctrl (P) | $3.81 \times 10^{-7}$ |
| *Pedobacter* | Ctrl (P) | $1.51 \times 10^{-5}$ |
| *Duganella* | Ctrl (P) | $5.15 \times 10^{-4}$ |
| *Pseudomonas* | Ctrl (P) | $5.16 \times 10^{-3}$ |
| *Listeria* | *L. moncytogenes* | $1.39 \times 10^{-10}$ |
| *Escherichia-Shigella* | *E. coli* O157:H7 | $6.97 \times 10^{-10}$ |
| *Faecalibacterium* | *E. coli* O157:H7 | $3.56 \times 10^{-6}$ |
| *Bacteroides* | *E. coli* O157:H7 | $4.34 \times 10^{-5}$ |
| *Eubacterium* | *E. coli* O157:H7 | $2.57 \times 10^{-3}$ |
| *Ruminococcus* | *E. coli* O157:H7 | 0.019 |
| *Sanguibacter* | *E. coli* O157:H7 | 0.032 |
| *Peredibacter* | *S.* infantis | $3.37 \times 10^{-3}$ |

Ctrl (C) represents the non-contaminated samples under the contamination groups; Ctrl (P) stands for the non-pathogenic samples under the pathogen groups.

**Supplementary Table 4.** *P* values of the biomarkers identified by the ANCOM-BC analysis for fresh produce in different quality groups.

| Biomarker | Group | *P* value |
|---|---|---|
| *Sphigomonas* | GQ | $4.54 \times 10^{-9}$ |
| *Pedobacter* | GQ | $3.16 \times 10^{-4}$ |
| *Parablastomonas* | GQ | $2.31 \times 10^{-3}$ |
| *Paracoccus* | GQ | $7.09 \times 10^{-3}$ |
| *Pir4_lineage* | GQ | 0.010 |
| *Nocardioides* | GQ | 0.048 |
| *Leuconostoc* | DQ | $8.23 \times 10^{-9}$ |
| *Gluconobacter* | DQ | $1.38 \times 10^{-7}$ |
| *Lactobacillus* | DQ | $1.16 \times 10^{-4}$ |
| *Acetobacter* | DQ | $5.8 \times 10^{-4}$ |
| *Clostridium* | DQ | 0.015 |

GQ represents good quality; DQ means decreasing quality

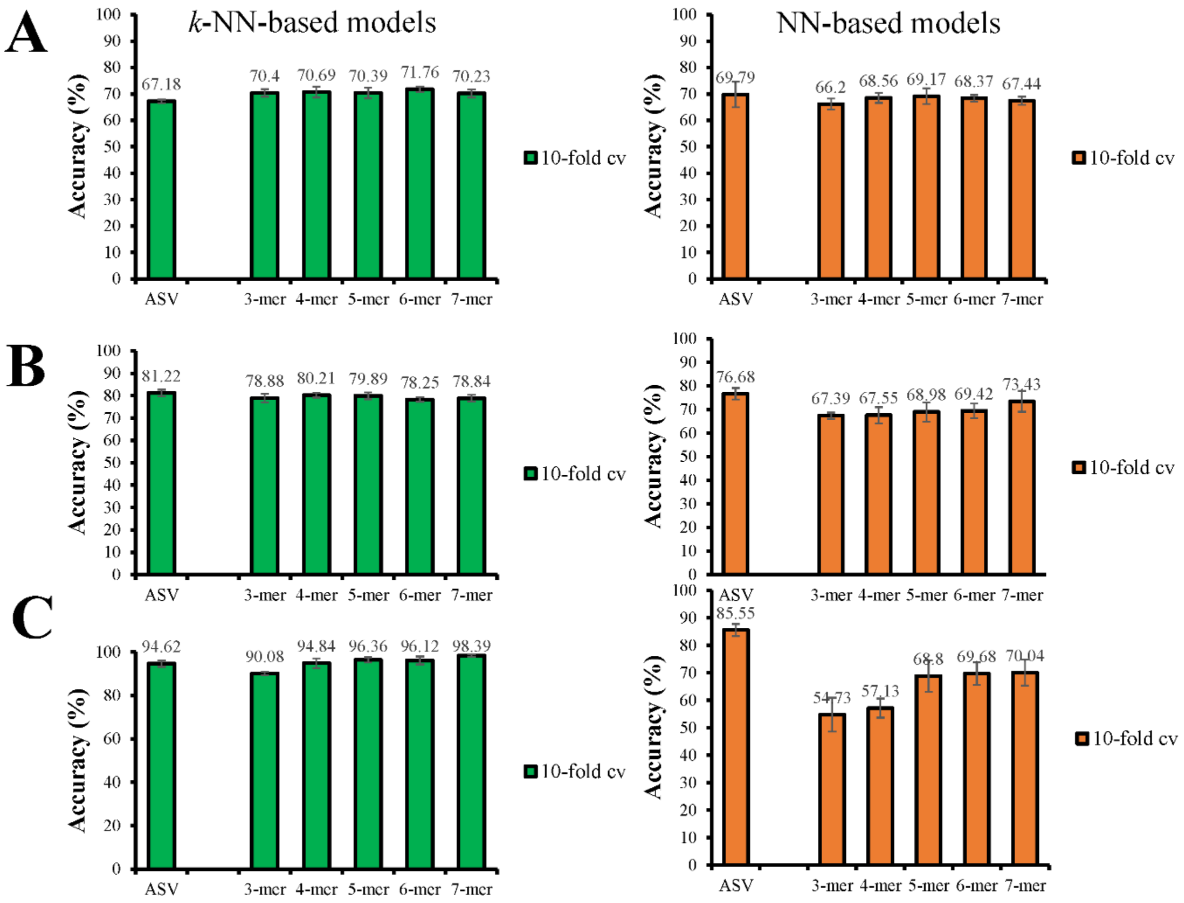


**Supplementary Fig. 1 | Comparison of the classification performance of PS classifiers based on *k*-NN and NN by using ASV datasets and hash datasets from 3-mer to 7-mer. (A)** Accuracy of PS classifiers based on *k*-NN and NN using ASV and 3-mer to 7-mer hash datasets of Zhang18. The trained models were validated by using the 10-fold CV method. **(B)** Accuracy of PS classifiers based on *k*-NN and NN using ASV and 3-mer to 7-mer hash datasets of LiaoRl21 (Liao and Wang, 2022; this study). **(C)** Accuracy of PS classifiers based on *k*-NN and NN using ASV and 3-mer to 7-mer hash datasets of LiaoSm21.

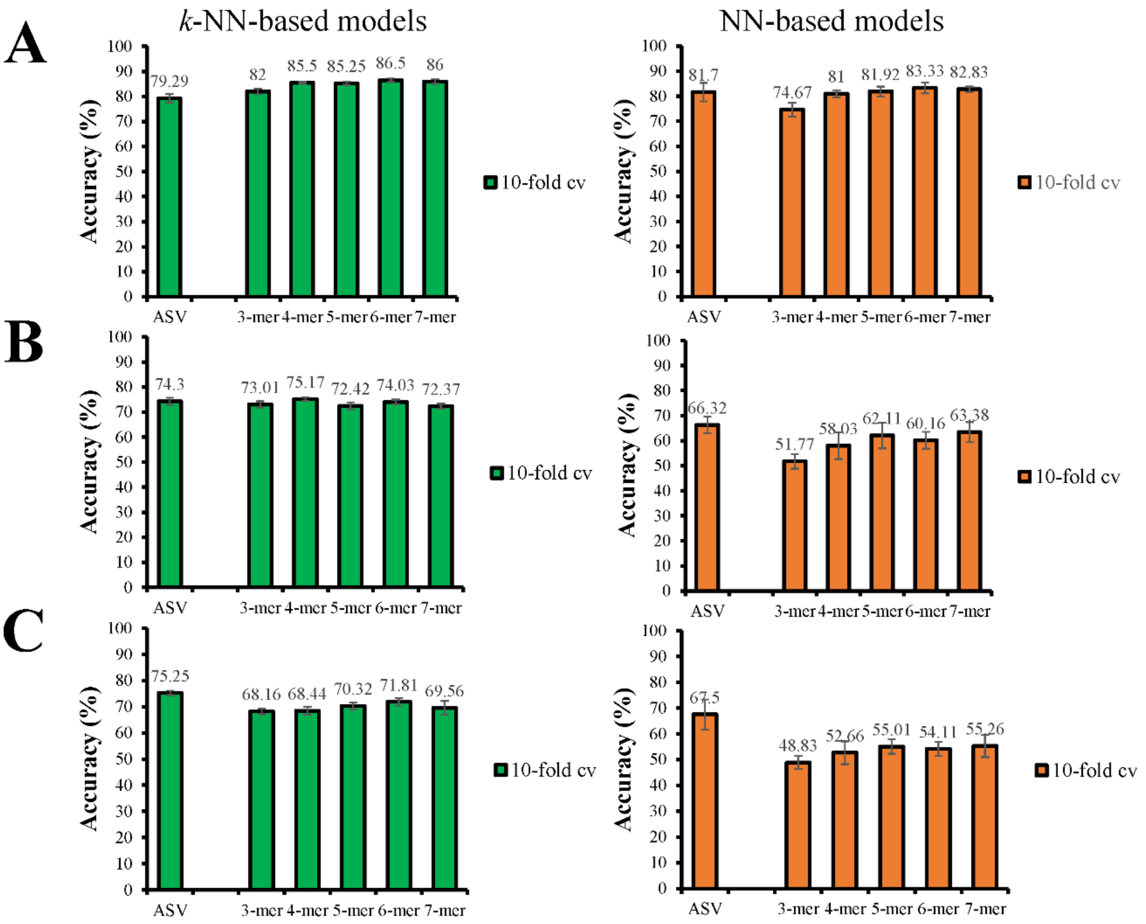


**Supplementary Fig. 2 | Comparison of the classification performance of PQ classifiers based on *k*-NN and NN by using ASV datasets and hash datasets from 3-mer to 7-mer. (A)** Accuracy of PQ classifiers based on *k*-NN and NN using ASV and 3-mer to 7-mer hash datasets of Kusstatcher19 project. The trained models were validated by using 10-fold CV method. **(B)** Accuracy of PQ classifiers based on *k*-NN and NN using ASV and 3-mer to 7-mer hash datasets of LiaoRl21 project (this study). **(C)** Accuracy of PQ classifiers based on *k*-NN and NN using ASV and 3-mer to 7-mer hash datasets of LiaoSm21 project.

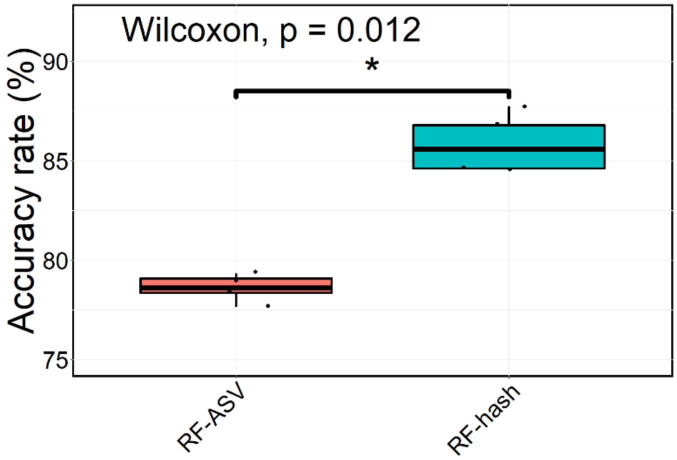


**Supplementary Fig. 3 | Accuracy of RF-based PS classifiers using integrated ASV and 7-mer hash datasets based on pathogen label.** Accuracy of PS classifiers using an integrated ASV and 7-mer hash datasets associated with PS (Zhang18, LiaoRl21, and LiaoSm21). The Wilcoxon rank sum test was used for the pairwise comparison of PS and PQ models.

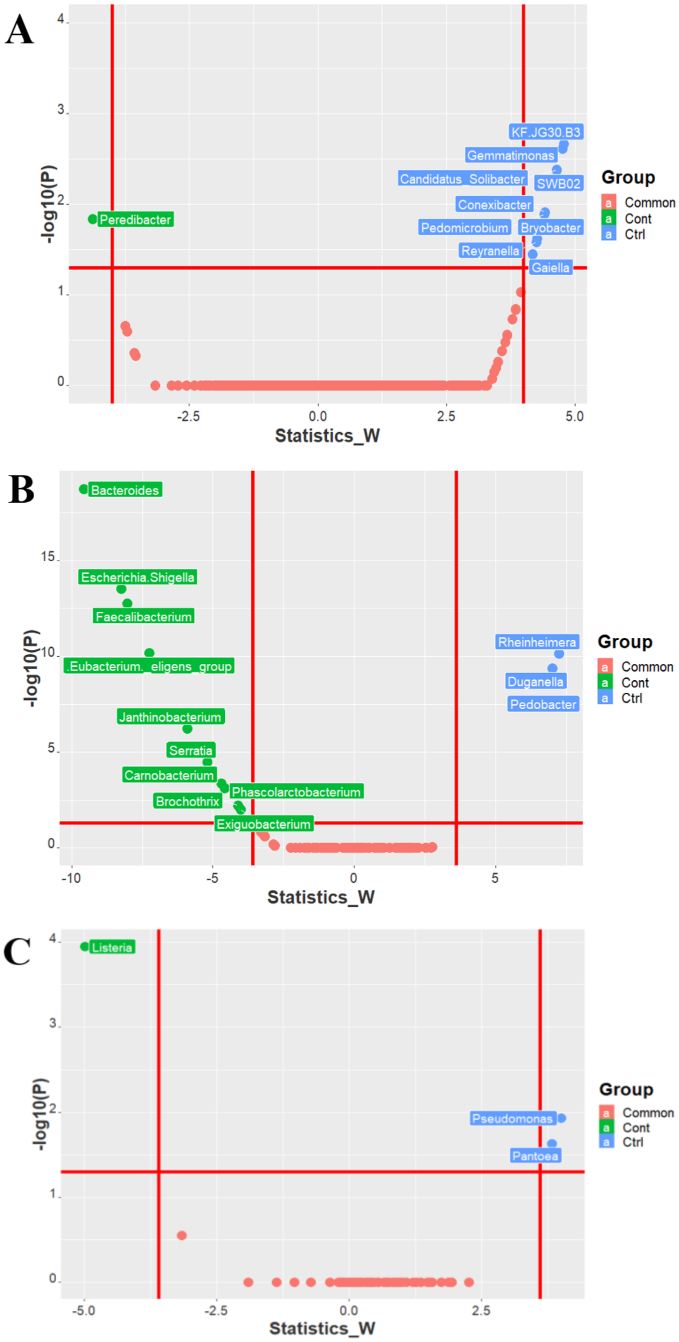


**Supplementary Fig. 4 | Taxonomic differential abundance analysis of individual fresh produce microbiome dataset related to PS.** (A) Volcano plot of the significantly differential abundance of identified bacteria at the genus level in different groups of samples from Zhang18. (B) Volcano plot of the significantly differential abundance of identified bacteria at the genus level in different groups of samples from LiaoSm21. (C) Volcano plot of significantly differential abundance of identified bacteria at the genus level in samples from LiaoRl21. Common represents bacteria with no significantly different abundance between non-contamination group (Ctrl) and contamination group (Cont).

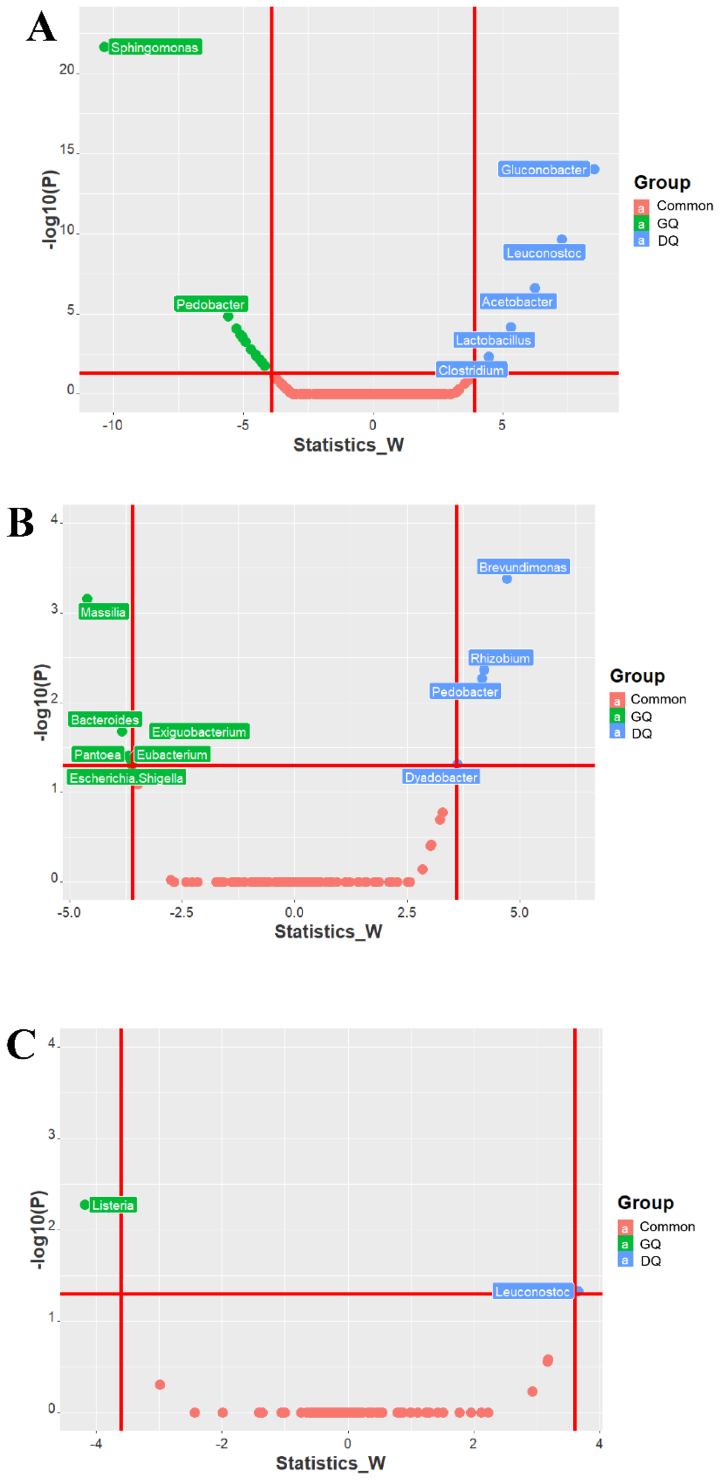


**Supplementary Fig. 5 | Taxonomic differential abundance analysis of individual fresh produce microbiome dataset related to PQ.** (A) Volcano plot of significantly differential abundance of identified bacteria at the genus level in different group of samples from Kusstatscher_2019. (B) Volcano plot of significantly differential abundance of identified bacteria at the genus level in different groups of samples from LiaoSm21. (C) Volcano plot of significantly differential abundance of identified bacteria at the genus level in samples from LiaoRl21. Common represents bacteria with no significantly different abundance between good-quality group (GQ) and decreasing-quality group (DQ).

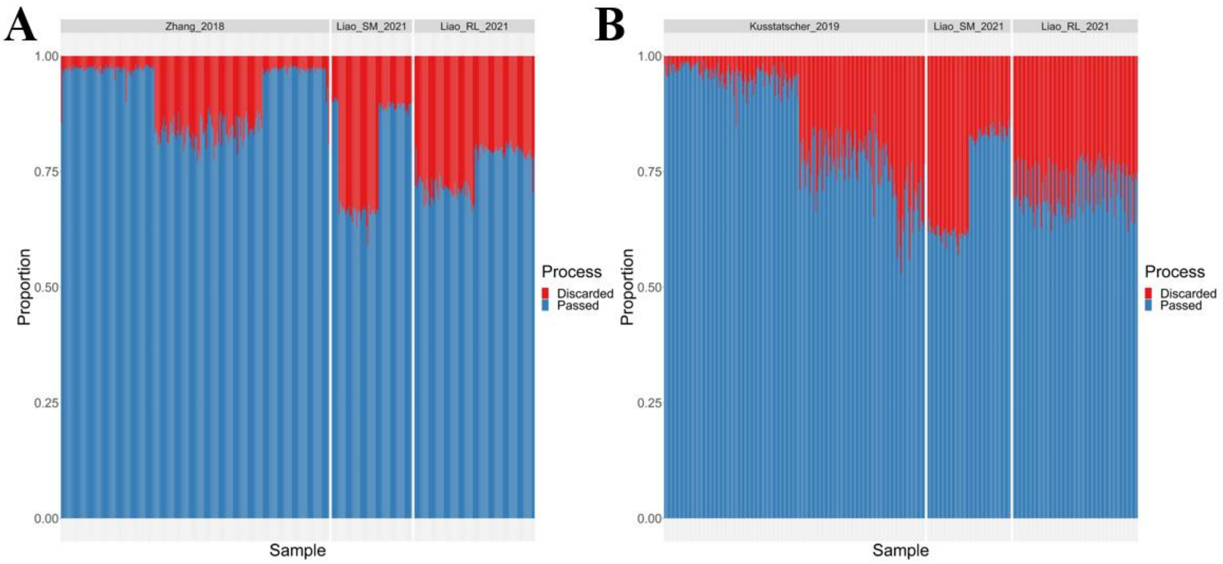


**Supplementary Fig. 6 | Proportion of sequence discarded from 16S rRNA sequence datasets of fresh produce microbiota related to PS and PQ during the denoising step by using the DADA2 plugin in QIIME 2.** (A) Sequence loss from three microbiome sequence datasets associated with PS, including projects Zhang18, LiaoSm21, and LiaoRl21. (B) sequence loss from three microbiome sequence datasets associated with PQ, including projects Kusstatscher19, LiaoSm21, and LiaoRl21.

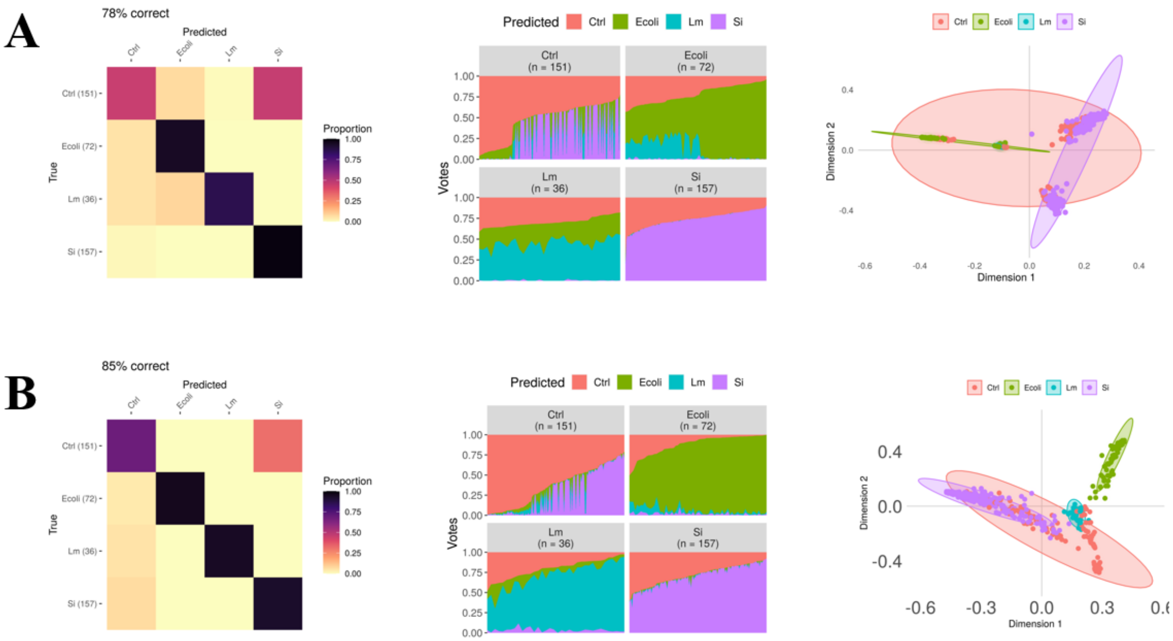


**Supplementary Fig. 7 | Plots of the confusion matrix (left), classification votes (middle), and MDS (right) generated from the RF-based models by using integrated ASV and 7-mer hash datasets associated with PS based on pathogen labels.** (A) RF-based models established by using integrated ASV datasets. (B) RF-based models established by using integrated 7-mer hash datasets. The circles with color around the sample dots on MDS plots indicate the classification to Ctrl: control samples (red), Ecoli: *E. coli* O157 contaminated samples (green), Lm: *L. monocytogenes* contaminated samples, and Si: *Salmonella* Infantis contaminated samples (purple).
